# Viral Packaging ATPases Utilize a Glutamate Switch to Couple ATPase Activity and DNA Translocation

**DOI:** 10.1101/2020.12.01.406595

**Authors:** Joshua Pajak, Rockney Atz, Brendan J. Hilbert, Marc C. Morais, Brian A. Kelch, Paul Jardine, Gaurav Arya

**Affiliations:** Dept. of Mechanical Engineering and Materials Science, Duke University, Durham, NC 27708; Dept. of Diagnostic and Biological Sciences, University of Minnesota, Minneapolis, MN 55455; Dept. of Biochemistry and Molecular Pharmacology, University of Massachusetts Medical School, Worcester, MA 01605; Dept. of Biochemistry and Molecular Biology, University of Texas Medical Branch, Galveston, TX 77550

**Keywords:** ASCE, ATPase, bacteriophage, crystal structure, DNA, DNA packaging, molecular dynamics simulations, molecular motor, mutual information

## Abstract

Many viruses utilize ringed packaging ATPases to translocate double-stranded DNA into procapsids during replication. A critical step in the mechanochemical cycle of such ATPases is ATP binding, which causes a subunit within the motor to grip DNA tightly. Here, we probe the underlying molecular mechanism by which ATP binding is coupled to DNA gripping and show that a glutamate switch residue found in AAA+ enzymes is central to this coupling in viral packaging ATPases. Using free energy landscapes computed through molecular dynamics simulations, we determined the stable conformational state of the ATPase active site in apo, ATP-bound, and ADP-bound states. Our results show that the catalytic glutamate residue transitions from an inactive to an active pose upon ATP binding, and that a residue assigned as the glutamate switch is necessary for regulating the transition. Further, we identified *via* mutual information analyses the intramolecular signaling pathway mediated by the glutamate switch that is responsible for coupling ATP binding to conformational transitions of DNA-gripping motifs. We corroborated these predictions with both structural and functional experimental data. Specifically, we showed that the crystal structure of the ADP-bound P74-26 packaging ATPase is consistent with the predicted structural coupling from simulations, and we further showed that disrupting the predicted signaling pathway indeed decouples ATPase activity from DNA translocation activity in the φ29 DNA packaging motor. Our work thus establishes a signaling pathway in viral DNA packaging motors that ensures coordination between chemical and mechanical events involved in viral DNA packaging.

## Introduction

Many double-stranded DNA viruses use powerful ATPase motors to package their chromosome into preformed viral capsids (1–4). These motors convert energy from ATP binding and hydrolysis into mechanical work to overcome large internal forces that oppose DNA confinement. Viral DNA packaging ATPases have two primary domains: the N-terminal domain (ATPase domain) and the C-terminal domain (which performs endonuclease activity in some viral packaging systems) (2). Based on sequence alignments of the ATPase domain, viral packaging ATPases are classified as members of the Additional Strand, Conserved Glutamate (ASCE) superfamily of P-loop NTPases (5). Other related ASCE families include AAA+ (ATPases Associated with diverse cellular Activities) and ABC (ATP Binding Cassette) transporter proteins (6, 7). A common feature among ASCE enzymes is the presence of a P-loop/Walker A (WA) and Walker B (WB) motifs which bind ATP and catalyze hydrolysis, respectively (8, 9). The feature that distinguishes ASCE enzymes from other superfamilies of ATPases is the presence of a conserved glutamate residue downstream of the WB motif that is necessary for catalytic activity (6, 10, 11). Like most ASCE systems, viral packaging ATPases assemble as oligomeric rings that thread their biopolymer substrate through their central pores (12, 13). Recently solved pentameric ring complexes of viral ATPases from the asccφ28 and φ29 bacteriophages show that the ATPase active site is located at the inter-monomer (or inter-subunit) interface (14, 15). Residues donated *in trans* promote events such as catalysis and nucleotide exchange, thus ensuring coordinated activity around the ATPase ring.

Coordinated activity of ATPase subunits has been extensively studied using single-molecule optical tweezers experiments (16). A key finding of these experiments is that all five subunits bind ATP prior to sequential and ordinal hydrolysis. Thus, the subunits must first bind ATP in a catalytically incompetent pose so as not to hydrolyze out of turn. There are at least two possible ways this can occur: 1) either the phosphates of the ATP substrate are misaligned upon initial binding and later align with catalytic residues to trigger hydrolysis, or 2) one or more catalytic residues of the active site are misaligned upon initial binding and later align to catalyze hydrolysis. The second of these alternatives is supported by the experimental observation that ATP-bound subunits grip DNA tighter than ADP-bound subunits (11, 17, 18). From this it can be inferred that the γ-phosphate of ATP is properly positioned to actuate a DNA-gripping signal, but the active site is not aligned to catalyze hydrolysis. In further support of the second alternative, we have recently proposed a cyclic helical-to-planar model of DNA translocation whereby *trans*-acting residues are misaligned for hydrolysis in the ATP-bound helical conformation of the ATPase ring (14). As subunits transition to the planar ring conformation during DNA translocation, these *trans*-acting residues align one interface at a time to catalyze hydrolysis, resulting in the ~2.5 bp DNA translocation substeps observed in single-molecule experiments.

The combination of a catalytically incompetent ATP-binding pose and simultaneous dependence on the γ-phosphate to actuate substrate gripping is not unique to viral packaging ATPases among the ASCE superfamily. This combination is also found in the AAA+ subfamily of ASCE enzymes. AAA+ motors coordinate these two tasks with an intramolecular mechanism known as the “glutamate switch.” In these systems, the glutamate switch is either a positively-charged Arg/Lys or a polar Gln/Asn/Thr/Ser, and is proposed to thwart ATP hydrolysis by holding the catalytic glutamate residue in an inactive pose until an external signal is received (19).

A key discriminator between the inactive and active poses are the χ_1_ and χ_2_ dihedral angles of the catalytic glutamate sidechain. If the (χ_1_, χ_2_) ordered pair point the catalytic glutamate’s carboxylate group towards ATP, then the pose is considered to be active. Conversely, if the (χ_1_, χ_2_) ordered pair point the catalytic glutamate’s carboxylate group towards the glutamate switch residue, the pose is considered to be inactive (19). A survey of AAA+ protein PDB structures in ATP- and ADP-bound states revealed tight clustering of (χ_1_, χ_2_) ordered pairs around the active and inactive poses, principally separated by an approximate 120o difference in χ_2_ (**Fig. 1**). Further, mutagenesis studies that targeted the glutamate switch residue showed that the glutamate switch participates in a signaling pathway between the ATPase active site and biopolymer substrate gripping residues. When the glutamate switch is mutated to an alanine residue, substrate-binding no longer upregulates ATPase activity (20, 21).

**Figure 1:**
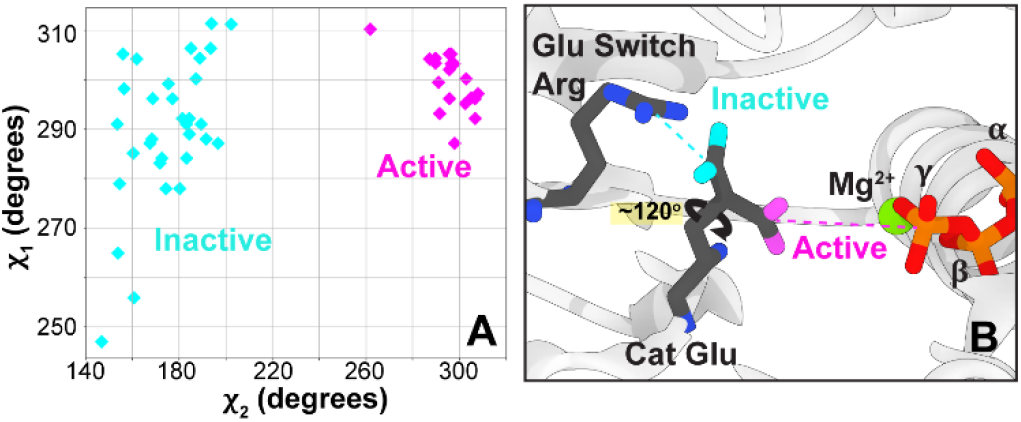
Schematic of glutamate switch. **A.** PDB structures of ATPases in ATP- and ADP-bound states show that the catalytic glutamate residue side chain dihedrals can be clustered and classified into “inactive” or “active” poses. Data reproduced from Zhang & Wigley (18) **B.** Depiction of these two poses. In the inactive pose, the catalytic glutamate’s carboxylate group points away from ATP and towards a glutamate switch residue; in the active pose, the catalytic glutamate’s carboxylate group points towards ATP to help catalyze hydrolysis. The phosphates of ATP are labeled, and Mg2+ is depicted as a green sphere.

Here we show that viral DNA packaging ATPases have adapted a glutamate switch residue as part of a signaling pathway that tightly couples ATP-binding to DNA gripping and ATP hydrolysis to DNA release. This coupling thus enables the overall coordinated activity of the motor necessary for efficient, processive translocation of DNA.

## Results

### Study design

To establish that viral DNA packaging ATPases utilize a glutamate switch to couple ATP-binding and DNA-gripping, it is necessary to show that:

1. The catalytic glutamate residue can be stable in either the “active” or “inactive” rotamer state.
2. Stability of the catalytic glutamate in the “inactive” state can be attributed primarily to a single residue (namely, the glutamate switch).
3. The glutamate switch can relay a bound-ATP signal to DNA-gripping residues.

To demonstrate **1**, we carried out atomistic molecular dynamics (MD) simulations of four different packaging ATPases and calculated the two-dimensional free-energy landscapes of their putative catalytic glutamate with respect to its (χ_1_, χ_2_) rotamer dihedral angles in different nucleotide-bound states. We assumed that if a glutamate switch were present, we would observe computed free-energy minima corresponding roughly to the “active” *and “inactive” clusters seen* experimentally (**Fig. 1**). To demonstrate **2**, we generated *in silico* mutants of the ATPases targeting the putative glutamate computed the free-energy landscapes. From how switch and re-computed the free-energy landscapes. From how the landscapes changed with respect to the type of mutation, we could infer the ability of the wild-type residue to function as a glutamate switch. To rule out that the tight-binding transition positions multiple residues that each contribute a little energy towards the cumulative catalytic glutamate free-energy landscape, we also considered ATPases held in a “pre-tight-binding” pose. To demonstrate **3**, we calculated mutual information from microsecond long MD simulations. Mutual information (MI) is a metric that quantifies correlation between two parties, in this case protein residues. From this correlation, we infer communication. If the glutamate switch mediates the signaling pathway, we would expect the glutamate switch to have high MI – and thus high communication – with both the ATPase active site and DNA gripping residues.

To validate our computational predictions, we sought both structural and functional characterization. Our simulations predict that a helical motif between the ATPase active site and DNA-gripping residues shifts in the presence of ATP, but not ADP. Thus, we solved the structure of the P74-26 packaging ATPase bound with ADP to compare against the previously reported ATP-analog bound structure (11). To interrogate the signaling pathway in an actively packaging assembly, we simultaneously characterized the ATPase activity and DNA translocation activity of mutant φ29 ATPases. This allowed us to determine if a mutant introduced in the signaling pathway would decouple these two otherwise tightly-coupled activities.

### Nucleotide affects the rotameric state of the catalytic glutamate

Available crystal structures of the Sf6, φ29 and asccφ28 packaging ATPases show that the predicted ASCE catalytic glutamate residues point their carboxylate group away from the ATPase active site and towards analogous arginine residues (**Fig. S1**). This is observed in the Sf6 crystal structures solved with either ATP or ATP-analog. Because this type of interaction is indicative of a glutamate switch mechanism in AAA+ proteins (**Fig. 1**), the crystal structures are the first indication that these arginine residues may act as glutamate switches.

Free-energy landscapes calculated for the proteins in the apo state show that the catalytic glutamate has two free energy minima, which correspond to the identified inactive and active rotamer states. In general, there is preference for the inactive pose (**Fig. S2**). This contrasts with the preference of an isolated glutamate residue in solution, which prefers the active pose (**Fig. S3**), suggesting that neighboring residues influence the catalytic glutamate’s preference for the inactive pose in the apo ATPases.

Free-energy landscapes calculated for the φ29 **(Fig. 2A,B)**, Sf6 **(Fig. 2C,D)**, and asccφ28 **(Fig. 2E,F)**ATPases in the ATP-bound and ADP-bound states show that the catalytic glutamate’s rotameric state varies with the presence or absence of the γ- phosphate of ATP. In the ATP-bound state, the catalytic glutamate prefers the active pose, pointing its carboxylate group towards ATP **(Fig. 2A,C,E)**. In the ADP-bound state (**Fig. 2B,D,F**), the global free-energy minimum of the catalytic glutamate changes to prefer the inactive pose, pointing its carboxylate group towards the putative glutamate switch arginines. This indicates that the presence of the γ-phosphate of ATP is necessary and sufficient to induce local conformational changes that stabilize the catalytic glutamate residue in the active pose. We note that the Walker A arginine residue moves into the binding pocket upon ATP-binding and moves out of the binding pocket in the absence of the γ-phosphate. We have previously reported that this conserved Walker A arginine in phage λ functions analogously to the “sensor II motif” arginine found in AAA+ enzymes, whose role is to sense the presence of the γ-phosphate and induce further conformational changes (10). The results presented here further support this assignment.

**Figure 2:**
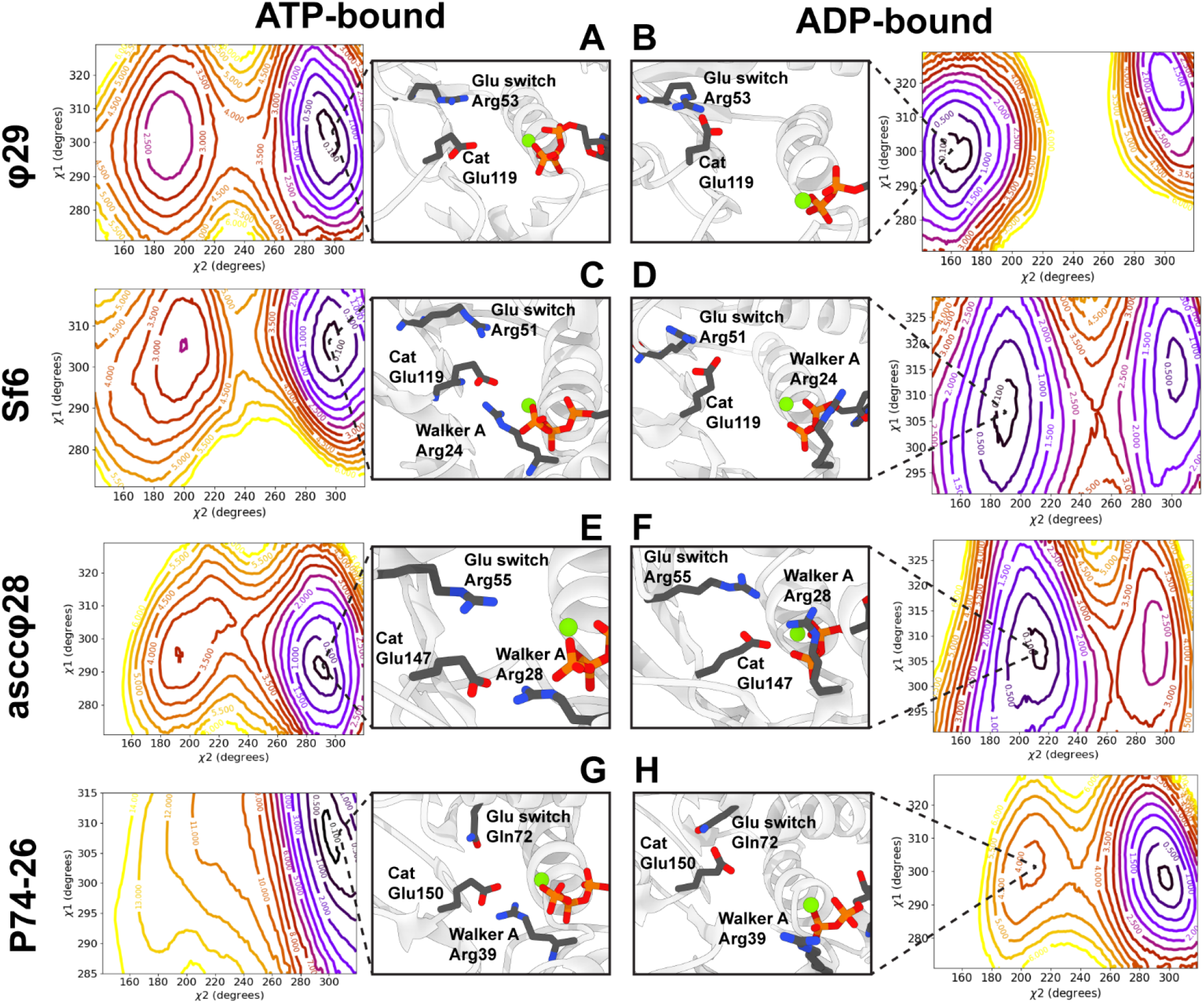
Viral DNA packaging ATPase catalytic glutamates pose is modulated by bound nucleotide. **(Left column)** Two-dimensional free-energy landscapes of the catalytic glutamate (χ_1_, χ_2_) dihedral angles from the φ29 **(A)**, Sf6 **(C)**, asccφ28 **(E)**, and P74-26 **(G)** packaging ATPases predict that the catalytic glutamate adopts an active pose upon ATP tight binding. Structures taken from MD simulations representing these poses are shown. For the φ29, Sf6, and asccφ28 ATPases, the inactive pose is metastable; for the P74-26 ATPase the inactive pose is unstable. **(Right column)** Free-energy landscapes of the catalytic glutamate (χ_1_, χ_2_) dihedral angles from the φ29 **(B)**, Sf6 **(D)**, and asccφ28 **(F)** packaging ATPases with bound ADP predict that upon phosphate release, the catalytic glutamate adopts an inactive pose and points towards a putative glutamate switch residue. In the case of P74-26 (H), while the active pose is still the preferred state, the inactive pose becomes metastable and thus shows response to absence of the ³-phosphate; this effect could be amplified from neighboring subunit interactions in the oligomeric state. Contour lines represent poses with constant free-energy in units of kcal/mol.

We also investigated the packaging ATPase from the P74-26 thermophilic phage. This ATPase does not contain an analogous glutamate switch arginine as in Sf6, φ29, and asccφ28 packaging ATPases described above. Instead, it contains an asparagine residue which we suspected could act as the glutamate switch due to its similar position relative to the catalytic glutamate, despite significant divergence in local tertiary structure (**Fig S4**). Indeed, the catalytic glutamate (χ_1_, χ_2_) free energy landscape in the P74-26 ATPase also varies with the presence or absence of the γ-phosphate of ATP (**Fig. 2G,H**). In the ATP-bound state, the catalytic glutamate is only stable in the active pose **(Fig. 2G)**. In the ADP-bound state, a metastability emerges in the inactive pose **(Fig. 2H)**. This metastability is also found in the apo state of the P74-26 ATPase (**Fig. S2**). While preference is not swapped between active and inactive like in the other three ATPases above, we note that this difference is perhaps expected because the P74-26 ATPase uses a polar residue for the glutamate switch as opposed to the positively charged residues in the φ29, Sf6, and asccφ28 ATPases. Thus, the nucleotide-bound state can be discriminated by the ATPase through the interaction between the catalytic glutamate and glutamate switch residues.

### A single residue is the glutamate switch

To determine the role of the ATP tight-binding transition on the rotameric state of the catalytic glutamate, we calculated the catalytic glutamate (χ_1_, χ_2_) free-energy landscapes of the φ29, Sf6, and P74-26 packaging ATPases in the “pre-tight-binding” conformations, where ATP is present but the ATPase active site has not yet adapted to its presence. These pre-tight-binding poses are characterized by the WA arginine pointing out of the binding pocket and an increased distance between the WA and WB motif backbones (**Fig. S5**), based on our two previous studies of tight-binding conformational changes (9, 10). We find that for φ29 **(Fig. 3A)**and Sf6 **(Fig. S6)**, the catalytic glutamate prefers the inactive pose prior to the ATP tight-binding transition. In the case of P74-26 **(Fig. 3C)**, the metastability of the catalytic glutamate’s inactive position is strengthened. However, it remained unclear whether the preference for the inactive pose of the catalytic glutamate could be attributed solely to its interaction with the putative glutamate switch residue, or if ATP tight binding induces multiple small conformational changes that cumulatively modulate the catalytic glutamate’s preference. Thus, we repeated the above calculations with catalytic glutamate mutant ATPases φ29 R53(L/K) and Sf6 R51(L/K). Leucine was chosen as a variant to remove the charge group from the putative glutamate switch residue while maintaining a similar bulk to the WT arginine. Lysine was chosen as a second variant in the hope that it would weaken, but not abrogate, its hold on the catalytic glutamate residue, so that the mechanism could be probed experimentally. Under the same conditions, where the WT catalytic glutamate prefers the inactive pose, the R→K mutants’ catalytic glutamate also preferred the inactive pose (**Fig. S6**). However, under these conditions the R→L mutants’ catalytic glutamate preferred the active pose (**Fig. 3B, Fig. S6**). Thus, the positive charge donated by the arginine residue is necessary to hold the catalytic glutamate in the inactive pose prior to the ATP tight-binding transition, and the arginine is thus assigned to be a glutamate switch residue.

Because the WT P74-26 packaging ATPase potentially uses a polar residue as its glutamate switch and the inactive pose is predicted to be metastable, it offers a unique opportunity to probe the effects of potentially strengthening, rather than weakening, the glutamate switch’s interactions with the catalytic residue. Thus, we chose to study the Q72K mutant to see if the change from polar to positively charged would cause the catalytic glutamate free-energy landscapes to be similar to other packaging ATPases. Indeed, the introduction of a positive charge in the P74-26 glutamate switch position strengthens the catalytic glutamate’s preference for the inactive pose compared to the WT (**Fig. 3D**). In total, our free-energy landscapes calculated for mutant ATPases enable us to assign these residues as glutamate switches.

**Figure 3:**
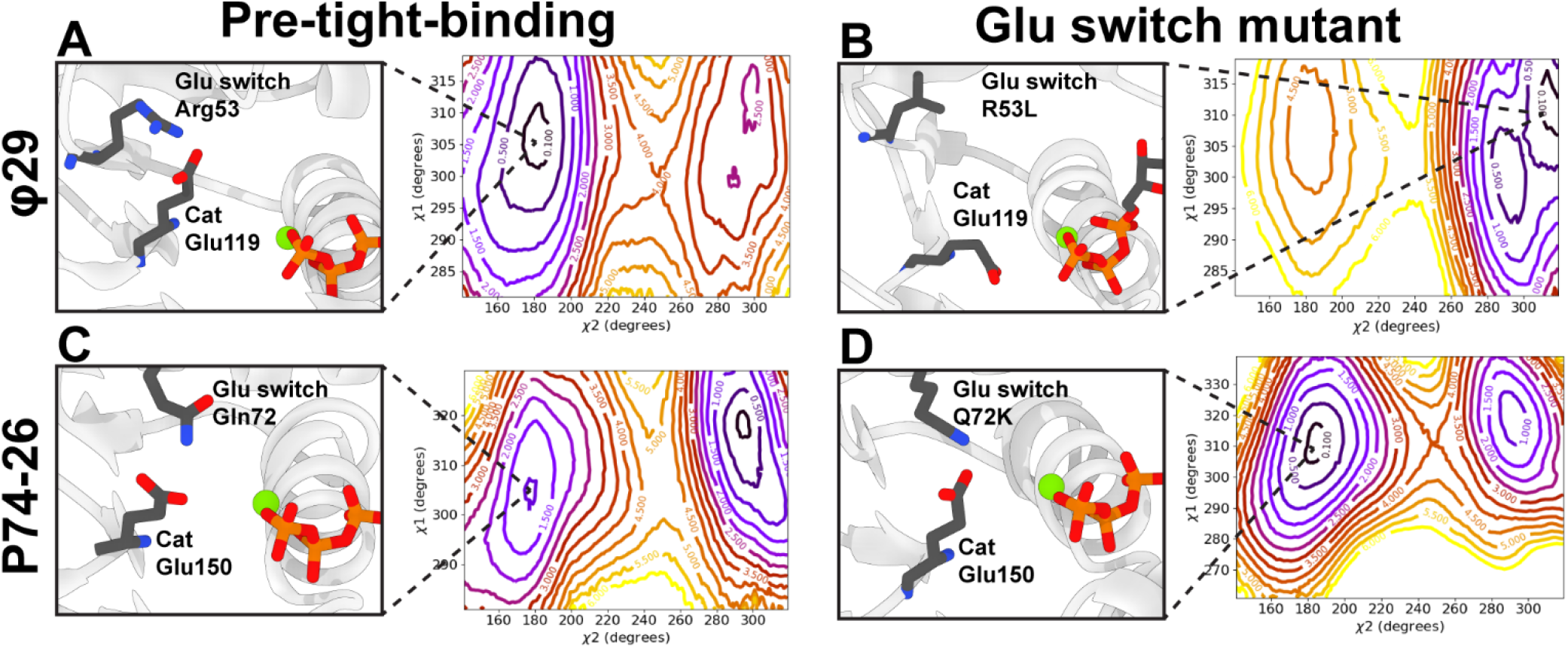
Point mutations are sufficient to nullify or amplify the glutamate switch behavior. **(Left column)** Free-energy landscapes of the catalytic glutamate’s (χ_1_, χ_2_) dihedral angles of wild-type ATPases in pre-tight-binding pose. For φ29 **(A)**, the inactive pose is preferred, and for P74-26 **(C)**, the inactive pose is metastable. **(Right column)** Free-energy landscapes of the catalytic glutamate’s (χ_1_, χ_2_) dihedral angles of glutamate switch mutants in pre-tight-binding poses. For φ29 **(B)**, the R53L mutant swaps preference from the inactive WT pose to the active pose, demonstrating that the charge carried by glutamate switch Arg53 is necessary to maintain inactivity prior to the tight-binding transition. On the other hand, P74-26 mutant Q72K **(C)** turns the metastable inactive pose into a globally stable pose, showing that adding positive charge to the polar glutamate switch Gln72 can amplify the hold of the glutamate switch on the catalytic glutamate. Thus, φ29 Arg53 and P74-26 Gln72 are both identified as glutamate switch residues. All contour lines represent changes in free-energy in units of kcal/mol. See Fig. S6 for φ29 R53K, Sf6 WT pre-tight-binding, Sf6 R51L, and Sf6 R51K free-energy landscapes.

### ATP binding triggers DNA gripping through a helical motif

To probe the relationship between ATP tight-binding and DNA interaction, we performed microsecond-long equilibrium MD simulations of ATP- and ADP-bound ATPases. In all cases, a helical motif between the ATPase active site and proposed DNA-gripping residues rotates upon ATP binding (**Fig. 4**). In the recently-solved cryo-EM structure of the φ29 stalled during packaging (15), Lys56 directly contacts the DNA phosphates. Lys56 is part of the rotated helical motif and is only a few residues downstream of the glutamate switch Arg53. Our simulations predict that relative to the ADP-bound state, ATP binding moves Lys56 **(Fig. 4A)**such that it would be donated into the pore of the ATPase ring. This suggests that the response of the catalytic glutamate and glutamate switch to ATP-binding is coupled to key residues believed to participate in DNA binding. For both the Sf6 **(Fig. 4B)**and asccφ28 **(Fig. 4C)**ATPases, our simulations predict that shifting the helical motif also repositions nearby proposed DNA-gripping motifs in a neighboring beta-hairpin through steric interactions. Thus, shifting this helical domain signals nucleotide occupancy to DNA-gripping residues. We note that it has not been experimentally determined which residues of the Sf6 packaging ATPase grip DNA. However, there are similar backbone shifts in Sf6 Arg82 as in its analog P74-26 Arg101, which has been demonstrated to grip DNA (22).

**Figure 4:**
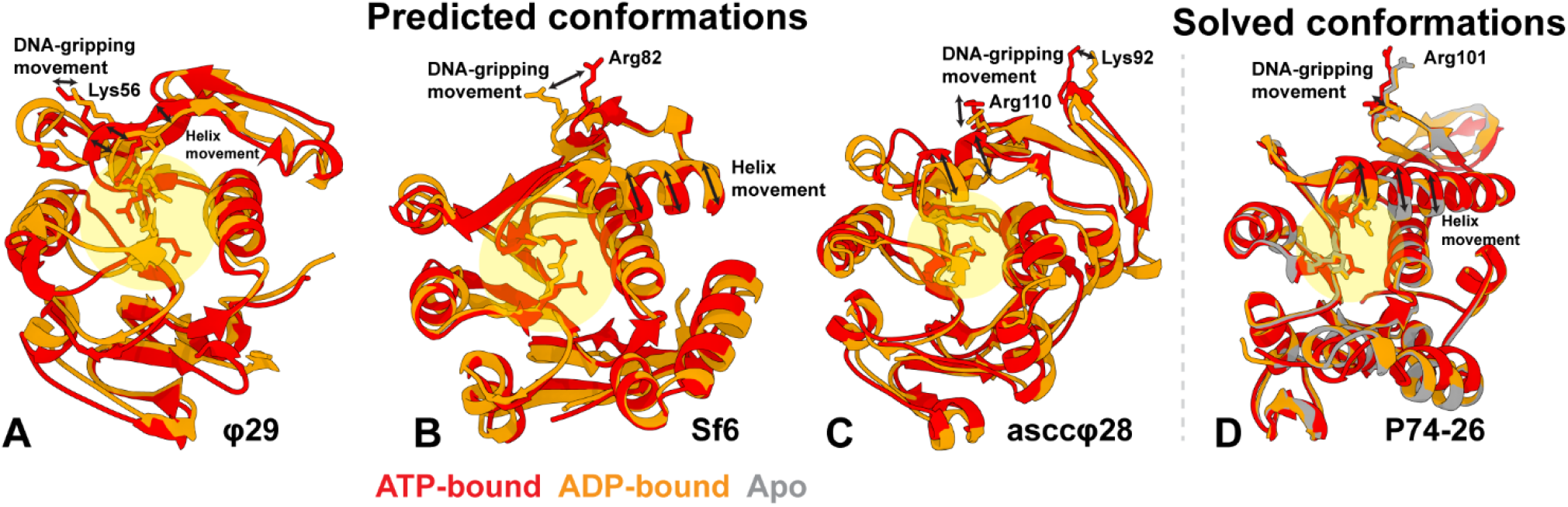
Displacements in a helical motif partitioning the ATPase active site and DNA-gripping residues in the ATP-bound conformation relative to the ADP-bound conformation. **(A-C)** MD simulations predict that a helical motif in the φ29 **(A)**, Sf6 **(B)**, and asccφ28 **(C)** ATPases in ATP-bound (red) and ADP-bound (orange) conformations moves in response to the γ-phosphate of ATP. **(D)** Experimentally determined structures of the P74-26 ATPase in ATP-bound (red), ADP-bound (orange), and apo (gray) conformations shows that this helical motif is only displaced in the ATP-bound state. In all cases, movement of this helical motif results in movement of DNA-gripping residues, suggesting a possible signaling pathway that promotes DNA-gripping upon ATP-binding. In all cases the catalytic glutamate and glutamate switch are shown as sticks and the active site is highlighted in yellow; DNA-gripping residues are also shown as sticks and labeled.

To validate these predictions, we determined the crystal structure of the ADP-bound P74-26 packaging ATPase to a resolution of |1.9 Å, using the same procedure used earlier for determining its apo and ATP-bound structures (11). Our structure of the P74-26 ATPase bound to ADP appears nearly identical to the apo state **(Fig. 4D)**; in contrast, the ATP-analog-bound state exhibits a pronounced displacement in the helical motif partitioning the ATPase active site from the DNA-gripping residues. Therefore, we infer that the presence of a γ-phosphate actuates the conformational change, in agreement with our computational predictions in the φ29, Sf6, and asccφ28 packaging ATPases. We note that the P74-26 ATPase is unique among the four ATPases studied here, as its glutamate switch is within this helical motif itself and not adjacent to the motif as in the φ29, Sf6, and asccφ28 ATPases; this difference in position may be related to the difference in the appearance of a polar residue as opposed to a positively charged residue in the glutamate switch position.

### Defining and testing the signaling pathway

To better understand the mechanism by which nucleotide occupancy is signaled through the helical motif and to the DNA-gripping residues, we carried out mutual information (MI) calculations. In particular, we calculated the MI between the rotamer/backbone dihedral angles of the identified glutamate switch residue and all other residues in each protein using the CARDS (Correlation of All Rotameric and Dynamical States) method (23). We found that in the simplest case of φ29 (where the primary DNA-gripping residue is located within the helical motif), the glutamate switch has high MI with both the catalytic glutamate and the predicted DNA-gripping motifs (**Fig. 5, Fig. S7**). In the other cases (where the primary DNA-gripping residue neighbors, but is not part of the helical motif), the glutamate switch has high MI with both the catalytic glutamate residue and the N-terminal of the rotated helical motif and the DNA-gripping residues have high MI with at least one residue in the helical motif. However, the glutamate switch and DNA-gripping residues do not have high MI with each other. We note that whereas CARDS can capture both direct MI (between residues *i* and *j*) and indirect MI (residue *i* interacting with residue *j* by way of residue *k*), it relies on correlating rotamer and backbone dihedral angles. Because of this dependence on dihedral angles, the target-site analysis may not necessarily predict high MI between two residues *i* and *j* if residue *i* induces a cartesian translation of *j* by changing backbone dihedral angles of residues *j* − 1 and *j* + 1. Thus, we assert that the two target-site analyses that predict that the glutamate switch and DNA-gripping residues both have high MI to different parts of the *same* helical motif still indicates communication between the two residues.

**Figure 5:**
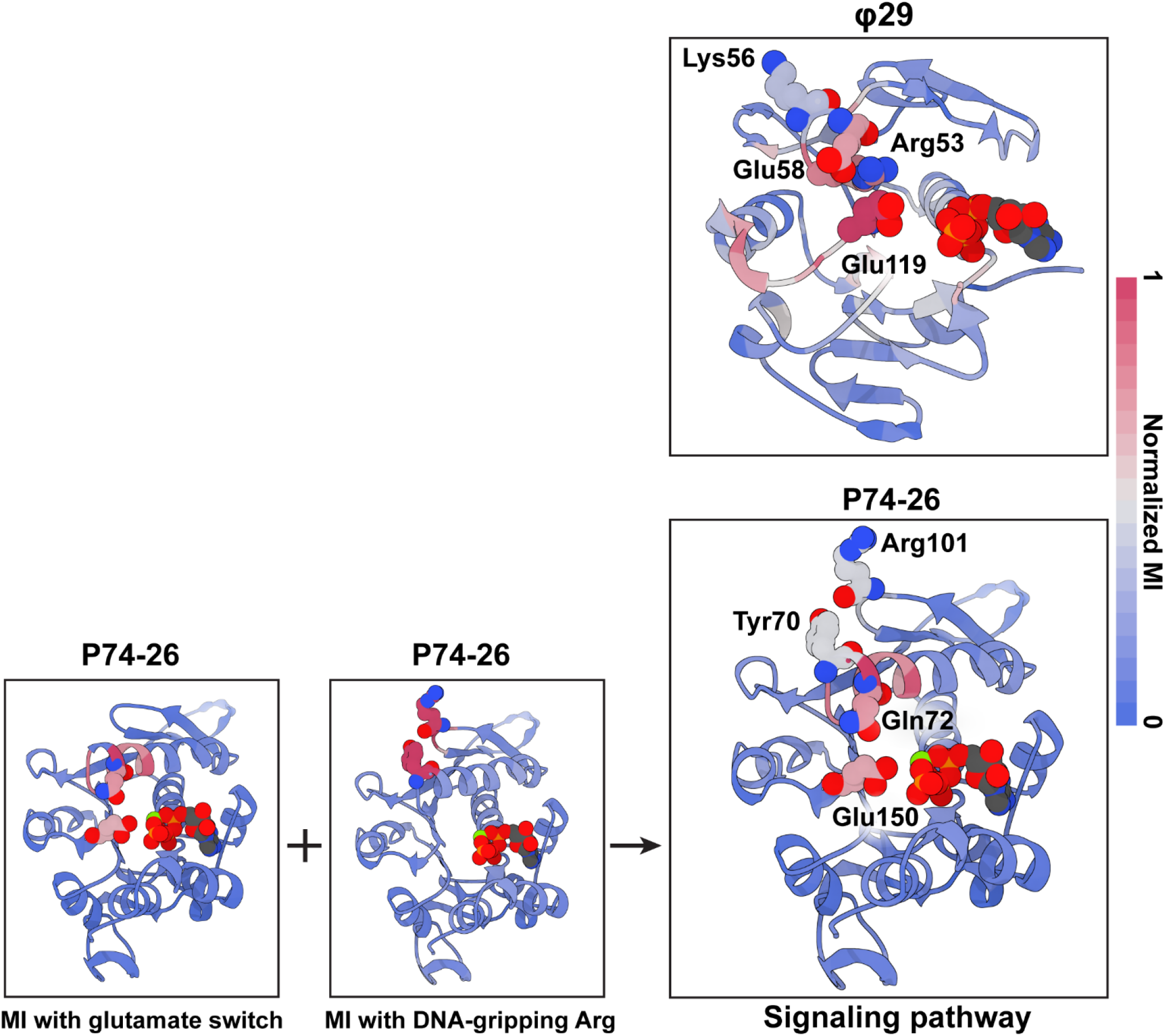
Mutual information calculated from MD simulations predicts a signaling pathway consistent with predicted and observed structure changes. **(Right)** Mutual information of each residue in the φ29 packaging ATPase calculated with respect to the glutamate switch Arg53. Red indicates high mutual information; blue indicates low mutual information. The glutamate switch has high mutual information with catalytic Glu119 and Glu58, which is part of the helical motif rotated during ATP binding (see **Fig. 4**). Lys56, which is experimentally known to grip DNA, is situated in the loop between Arg53 and Glu58. Thus, the catalytic Glu119 response to ATP/ADP can be signaled to DNA-gripping Lys56 by way of Arg53 and Glu58. (**Below**) Mutual information of each residue in the P74-26 ATPase calculated with respect to the glutamate switch Gln72. The glutamate switch has high mutual information with both the catalytic Glu150 and the helical motif rotated during ATP-binding (see **Fig. 4**). Mutual information of each residue with the DNA-gripping Arg101 shows high mutual information with a bulky residue in this same helical motif. Putting together these mutual information calculations, a signaling pathway connecting the ATPase active site and DNA-gripping residues is derived. Similar pathways can be derived for asccφ28 and Sf6 ATPases (see Fig. S7).

To validate our computational predictions, we experimentally characterized select mutant enzymes predicted to disrupt the signaling pathway. For this purpose, we chose the φ29 system, which is one of the most well-developed packaging systems among those studied. Importantly, assays for this system can be performed on motors that are actively packaging DNA into viral procapsids. This allows for separate and simultaneous measurements of ATPase activity and DNA translocation activity. Thus, it is an ideal system to study the coupling between the ATPase active site and DNA gripping residues. We found that changes to the glutamate switch residue itself either do not affect overall function (R53K) or abrogate function altogether (R53L), which may be due to slight changes in protein folding leading to defective oligomerization (**Fig. 6**). We also interrogated the signaling pathway without disturbing the glutamate switch residue. Our MI calculations predicts that the glutamate switch Arg53 interacts through Glu58 to regulate the position of DNA-gripping Lys56 (**Fig. 5**). Thus, we speculated that E58A would be a viable variant to disrupt the signaling pathway. Molecular dynamics simulations of an ATP-bound E58A variant predicts that its DNA-gripping Lys56 is positioned similarly to the apo state in WT (**Fig. 6**). Thus, we predicted that the E58A variant may not competently bind DNA despite not changing any DNA-gripping residue. We then separately characterized the ATPase activity and packaging activity of an E58A variant and found that while the ATPase activity of the E58A variant is at near-WT levels, the DNA translocation activity is drastically reduced. This is ostensibly because the motor can no longer grip DNA tightly in response to ATP-binding (**Fig. 6**). The experimentally observed decoupling of ATPase activity and DNA translocation supports the computational assignment that the glutamate switch signals DNA-gripping Lys56 through Glu58 in the φ29 packaging ATPase.

**Figure 6:**
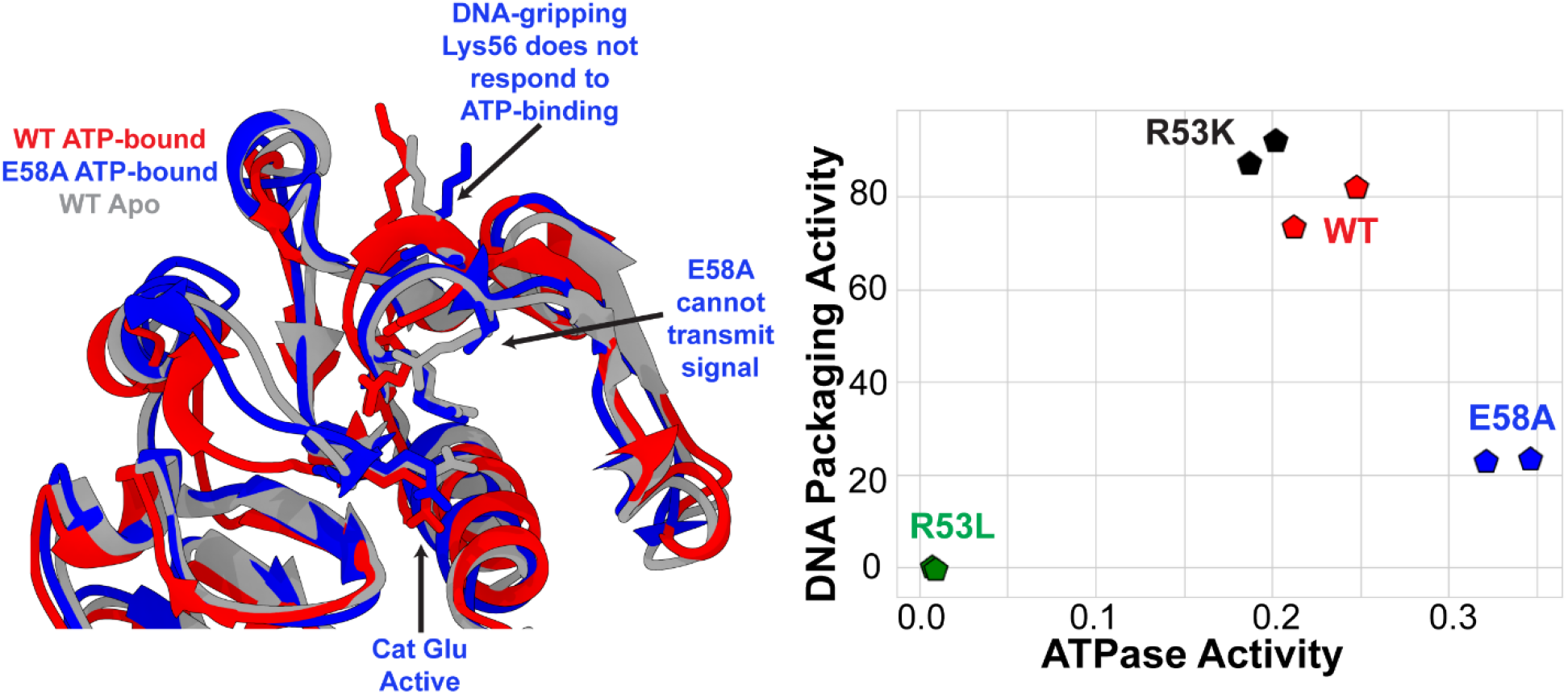
φ29 E58A mutant breaks the signaling pathway between ATPase active site and DNA-gripping Lys56. **(Left)** WT φ29 in ATP-bound (red) and apo (gray) conformations obtained from MD simulations predict that actuation of the catalytic Glu119 in response to ATP binding transmits a signal through Arg53 and Glu58 to displace Lys56 and better grip DNA (see **Fig. 5**). MD simulation of ATP-bound φ29 E58A mutant predicts that the DNA-gripping Lys56 is in a similar pose as the apo state, despite the actuation of the catalytic Glu119. This suggests that Glu58 is necessary to couple ATP-binding to DNA-gripping. **(Right)** Experimental ATPase and packaging assays of WT φ29 and select mutants. Glutamate switch mutant R53K is similar to the WT, whereas R53L abrogates both packaging and ATPase activity, likely due to folding or assembly defects. On the other hand, mutant E58A competently binds and hydrolyzes ATP but cannot package DNA, as would be expected if Glu58 is necessary to signal the ATP-binding event to DNA-gripping residues.

## Discussion

Recently, our work on determining atomic-resolution structures of the pentameric packaging ATPase complexes of asccφ28 and φ29 led to the derivation of a helical-to-planar ring model of viral DNA translocation (14, 15). A key tenet of this model is that ATP hydrolysis and subsequent product release in one subunit do not directly translocate DNA. Rather, product release causes the hydrolyzing subunit to lose its grip of DNA, which allows the neighboring ATP-bound subunit to rotate about its strained lid subdomain and drive DNA past the hydrolyzing subunit and into the capsid (**Fig. 7,**center panel). After the translocation event, planar alignment of the ATP-bound subunit with its now ADP-bound neighbor allows residues from the ADP-bound neighbor to be donated *in trans* to catalyze hydrolysis. This coordinated series of events then proceeds sequentially and ordinally around the ring.

**Figure 7:**
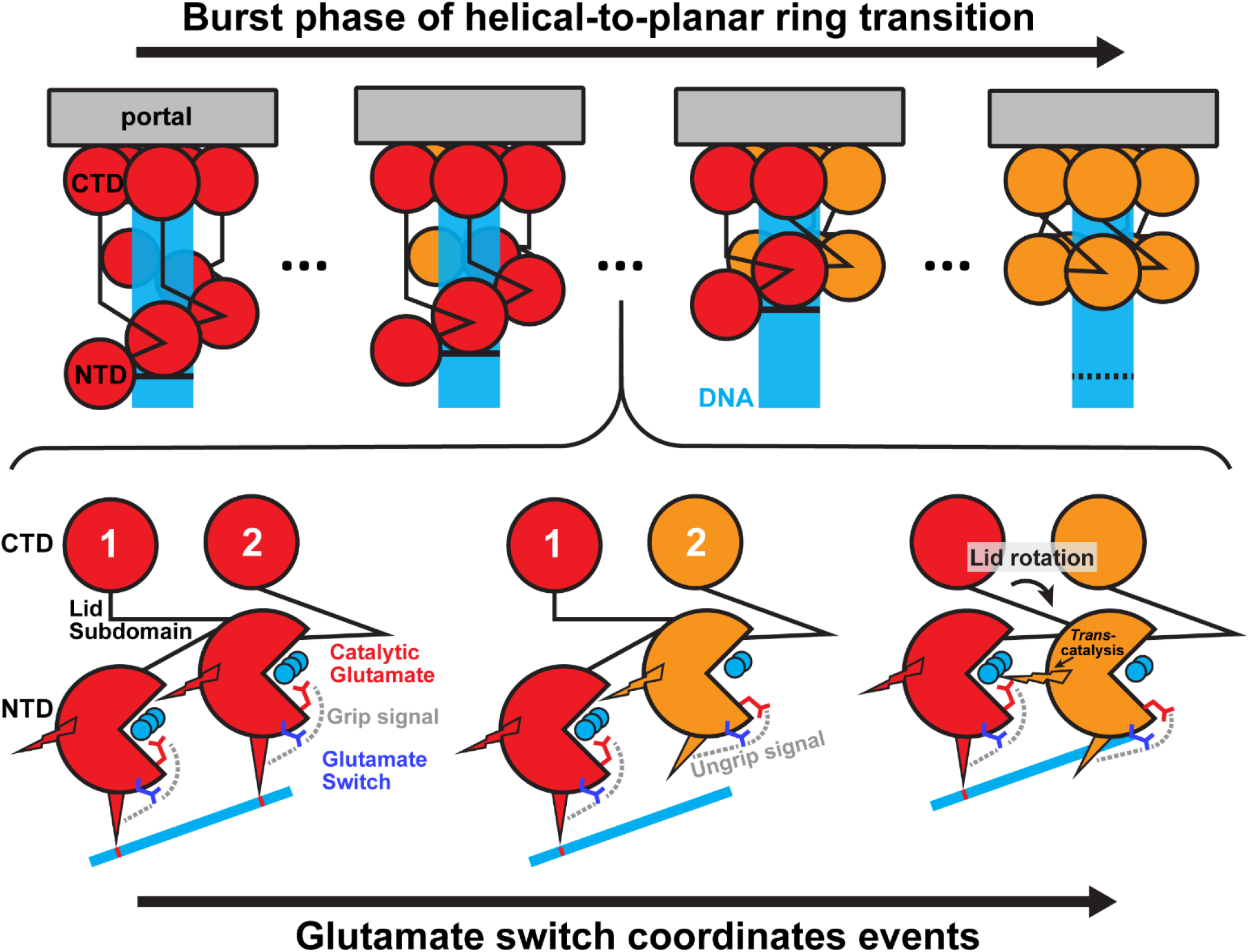
Glutamate switch coordinates sequential DNA packaging during the helical-to-planar ring transition burst phase. **(Top panel)** Simplified depiction of the helical-to-planar ring model proposed in Pajak et al. (14). Initially all five subunits are ATP bound (red) and their N-terminal ATPase domains are helically arranged. ATP hydrolysis and phosphate release causes the N-terminal ATPase domains, which are now all ADP bound (orange), to collapse to a planar arrangement, driving DNA by one helical turn into the capsid. Each subunit is shown as two circles representing the N- and C-terminal domains connected by a line representing the lid subdomain. DNA is shown as a transparent rectangle, and the contact between the lowest subunit and DNA is highlighted as a solid bar to guide the eye; in the end, the next repeat of DNA is represented with a dashed contact. **(Bottom panel)** Detailed depiction of events regulated by the glutamate switch. Two of the five subunits are highlighted. Initially in the helical arrangement, both subunits are ATP bound (red subunits; ATP shown as three cyan circles), which positions both catalytic glutamate residues in the active pose and signals DNA-gripping residues to grip DNA (signaling pathways are identified by labeled dashed lines). After hydrolysis in the upper subunit 2, its catalytic glutamate returns to the inactive pose and signals DNA release. When the ADP-bound (orange subunit; ADP shown as two cyan circles) subunit 2 releases DNA, the strained lid subdomain of subunit 1 rotates and brings its ATPase domain in plane with subunit 2. This translocates DNA by ~2.5 bp into the procapsid. Upon alignment, trans-acting residues (lightning bolt) donated from subunit 2 are able to catalyze hydrolysis in subunit 1. This mechanism propagates around the ring to ultimately translocate DNA ~10 bp, or one helical turn, into the procapsid.

The assertion that release of inorganic phosphate caused a subunit to lose grip of DNA is supported by experimental evidence showing that the ADP-bound state does not grip DNA as tightly as the ATP-bound state (11, 17, 18), and by computational predictions showing that the ADP-bound state does not donate positively-charged residues into the pore of the pentamer as well as the ATP-bound state (14). However, the exact mechanism that coupled the presence of the γ-phosphate of ATP to DNA binding was unclear. The glutamate switch mechanism detailed above fills this crucial gap in knowledge. Because the glutamate switch signaling pathway is coupled with conformational changes of the catalytic glutamate residue, it is uniquely positioned to respond to the presence and/or absence of the γ-phosphate of ATP. Further, consistent with glutamate switches found in AAA+ enzymes, the pathway is found to signal key substrate-gripping residues in all viral packaging ATPases examined here.

In the context of our helical-to-planar ring model (14, 15), the glutamate switch can send either an “on” or a “off” signal to the DNA-gripping residues **(Fig. 7)**. Upon ATP tight-binding, the catalytic glutamate residue points its carboxylate group towards the γ-phosphate of ATP and away from the glutamate switch residue. This sends the “on” signal to the DNA-gripping residues, moving them into the pore to tightly grip DNA. Although the *cis-*acting catalytic glutamate residue is properly aligned for hydrolysis at this point (subunit 1, **Fig. 7**), the *trans*-acting catalytic residues of the neighboring subunit (subunit 2, **Fig. 7**) are not aligned, and therefore subunit 1 holds ATP in a catalytically incompetent pose. After subunit 1 translocates DNA, subunit 2’s *trans*-acting catalytic residues align, and hydrolysis is catalyzed in subunit 1. Subsequent release of the inorganic phosphate from subunit 1 points its catalytic glutamate’s carboxylate group towards the glutamate switch residue. This interaction sends the “off” signal to subunit 1’s DNA-gripping residues, so that subunit 1 loses grip of DNA and the process repeats around the ring.

Our results demonstrate that, in viral DNA packaging ATPases, the primary function of the glutamate switch is to communicate ATP binding and hydrolysis to DNA-gripping residues and not to hold the catalytic glutamate in an inactive pose. This allows the motor to conform its quaternary structure to the helical substrate in a nucleotide-dependent manner. This raises the question as to the glutamate switch’s function in AAA+ motors. It was thought that the glutamate switch’s primarily role was to hold the catalytic glutamate inactive (19). Recently, many AAA+ motors have also been solved as helical rings that track their substrate in the ATP-bound state, with ADP-bound or apo subunits losing grip of substrate and not conforming to the overall helical pitch of the AAA+ ring (24–27). Thus, we posit that AAA+ motors may also utilize glutamate switches to ultimately translocate their substrate by modulating interaction between a subunit and substrate coupled to ATPase activity.

## Materials and Methods

### Structure preparation for simulations

Initial structures for packaging ATPases were taken from their respective PDB deposits: P74-26 (4ZNI, apo; 4ZNL, ATP-analog-bound) (11), Sf6 (4IDH apo; 4IEE, 4IFE, ATP-analog- and ATP-bound) (28), φ29 (5HD9, ATP-analog bound, ATP-analog not resolved) (13), and asccφ28 (14). Simulations of the ADP-bound P74-26 structure start from the solved crystal reported herein. Mutant proteins were generated *in silico* by replacing side chains with rotamers chosen from the Dunbrack rotamer library (29) via the *swapaa* command implemented in UCSF Chimera (30). Because the C-terminal domain does not directly interact with the ATPase active site during packaging, the C-terminal domains of the Sf6 and asccφ28 packaging ATPases were truncated to significantly reduce system size and computational cost; structures of P74-26 and φ29 only contain the ATPase domain.

### Simulation methodology

All-atom molecular dynamics (MD) simulations were performed with a 2 fs time step integrator in the AMBER16 or AMBER18 package using the Amber ff14SB force field (31) to describe protein interactions. The SHAKE algorithm was used to constrain bonds connecting hydrogen atoms to heavy atoms. Simulations were performed with GPU-accelerated sampling utilizing the particle mesh Ewald summation to correct for long-range interactions (32). All proteins were centered in an octahedral periodic box with minimum 14 Å of padding. The box was filled with explicit solvent using the TIP3P model for water. ATP parameters were taken from the AMBER parameter database (33). Each system was subjected to 300 steps of steepest descent and conjugate gradient energy minimization. Systems were heated slowly from 100 K to 310 K over the course of 100 ps in the canonical (NVT) ensemble. Then each system was equilibrated for 100 ns in the isobaric-isothermal (NPT) ensemble at 1 bar before free-energy sampling was performed. The Monte Carlo barostat and the Langevin thermostat were used to control pressure and temperature.

### Free-energy calculations

We used the combined approach of umbrella sampling (US) and the weighted histogram analysis method (WHAM) to calculate the two dimensional free-energy landscapes. The (χ_1_, χ_2_) angles of the catalytic glutamate residue were harmonically restrained with spring constant k = 30 kcal/mol/rad2 to windows spanning the range of the desired landscape. Because most of the distinction between active and inactive poses manifests as change in the χ_2_ variable, we separated χ_2_ windows by 7o, and χ_1_ windows by 20o to ensure ample sampling along the primary reaction coordinate. To hold an ATPase in the “pre-tight-binding” pose, the Walker A arginine side chain dihedral angles were restrained by the same harmonic potential. Each window was simulated for 3 ns and data points were collected every 100 fs. The potential of mean force was calculated using *wham-2d* (34), as described by Kumar et al.(35).

### Mutual information calculation

To predict signaling pathways connected to the glutamate switch, we calculated the mutual information (MI) of the side chain and backbone dihedral angles with every other residue in the protein. We used the Enspara package (36) to calculate the CARDS (Correlation of All Rotameric and Dynamical States; (23) from microsecond-long equilibrium simulations, saving frames every 25 ps. The holistic correlation *I_H_* between two dihedral angles X and Y is defined to be 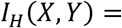 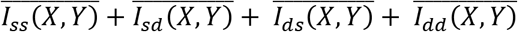 that is, CARDS calculates MI from concerted structural changes 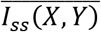 (e.g. a rotamer choice in one residue that influences a backbone rotamer choice in another), structural changes that coincide with conformational disorder 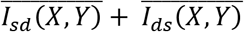 (e.g.a rotamer choice in one residue that allows another residue to become flexible), and conformational disorder 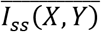 (e.g. flexibility in one residue correlating with flexibility in another residue). We then performed a target site analysis, which aggregates all the mutual information with respect to an inputted target site. The calculated target mutual information was normalized by the highest mutual information calculated for a single residue-residue pair and then input into PDB files as the temperature factor for visualization.

### Crystallization and structure determination

The P74-26 ATPase domain was expressed and purified as previously described (11). Concentrated protein was diluted to 5 mg/mL in 25 mM Tris (pH 8.5), 125 mM Sodium Chloride, 10% (v/v) Glycerol, 10 mM ADP, 10 mM Magnesium Chloride, and 10 mM DTT. ATPase domain crystals were grown using the hanging drop vapor diffusion method, with drops consisting of equal volumes of the protein solution and crystallization well solution (100 mM Sodium Acetate:HCl pH 5.0, 0.5 M ammonium sulfate). Crystals were harvested and flash frozen after a 30s soak in cryoprotectant containing ADP (50 mM Sodium Acetate:HCl pH 5.0, 0.8 M Ammonium Sulfate, 10 mM ADP, 10 mM Magnesium Chloride, and 30% Ethylene Glycol).

Crystallographic data were collected at the Advanced Light Source beamline 5.0.1. The data were processed with HKL3000 (37). Molecular replacement was accomplished with the SeMet model, as previously described (11). Molecular replacement and refinement were performed with Phenix (38) and model building with Coot (39). In addition to standard positional and B-factor refinement for the protein, we also directly refined the occupancy of the ADP molecule.

### ATPase and DNA translocation activity assays

DNA-packaging and ATPase assays were run simultaneously using a randomized, predetermined order of reactions of two independently made, replicate batches for each protein. φ29 proheads, DNA-gp3, and gp16 were prepared as previously described (40). The in vitro DNA packaging assay is based on a DNase protection assay (41). The ATPase assay uses Malachite green to measure the production of inorganic phosphate (42).

Purified gp16 was diluted to 100 ng/ml (2.5 μM) with SUMO protease cleavage buffer containing 50 mM Tris-HCl (pH 8), 400 mM NaCl, 10% v/v glycerol, and 3mM TCEP. sus8.5(900)-sus14(214)-sus16(300) proheads were diluted to 1 mg/ml (83nM) buffer containing 50 mM Tris (pH 7.8) and 10 mM MgCl_2_. φ29 DNA-gp3 was diluted to 250 ng/ml (21nM) with sterile dH_2_O.

Proheads and DNA-gp3 were combined in reaction buffer (100 mM Tris (pH 7.8), 20 mM MgCl_2_, and 20 mM NaCl) to final concentrations of 10 nM and 62.5 nM, respectively. Reaction mixtures were prepared by adding 7 μl of gp16 solution with 56 μl of the prohead/DNA-gp3 mixture, then incubated in ice for 15 minutes. Zero-time DNA packaging samples were made by combining 18 μl of the reaction mixture with 4 μl of a solution containing 5ng/ml DNase I, 50 mM Tris (pH 8), 0.05 mM CaCl_2_, 10 mM MgCl_2_, and 50% glycerol, incubating in a room-temperature water bath for 15 minutes, and then chilled in ice.

DNA-packaging and ATPase assays were conducted simultaneously from the same samples. The reactions were started by combining 40.5 μl of the above reaction mixture with 4.5 μl of 5 mM ATP. A zero-time ATPase sample was prepared by immediately removing 10 μl from the reaction, which was combined with 160 μl of Malachite green reagent. After 1 minute, the Malachite green reaction was quenched with 20 μl of 34% w/v Na-citrate and allowed to remain at room temperature. The remaining 35 μl of the reaction was placed in a room temperature water bath. At 15 min, a second ATPase sample was prepared by removing 10 μl from the reaction, which was combined with 160 μl of Malachite green reagent and processed as above. Simultaneously, 15 min DNA packaging samples were prepared by combining 20 μl of the reaction with 2 μl of solution containing 10 ng/ml DNase I, 50 mM Tris (pH 8), 0.05 mM CaCl2, 10 mM MgCl2, and 50% glycerol, incubated in a room-temperature water bath for 15 minutes, and then chilled in ice.

Zero-time and 15-minute time DNA-packaging samples were further processed by combining each sample with 2 μl of solution containing 0.25 M EDTA and 5 mg/ml proteinase K and incubating at 65°C for 30 minutes. The packaged, undigested DNA was resolved in an agarose gel and the densities of the packaged DNA and corresponding backgrounds were analyzed with Carestream Molecular Imaging Software 5.3.4.17821 (BioRad). 100 μl aliquots of zero-time and 15-minute time ATPase samples were transferred to 64-well plates and absorption were measured at 620 nm with a BioTek plate reader. Zero-time measurements were subtracted from their respective 15-minute time measurements and the ATPase measurements were plotted on the DNA-packaging measurements.

## Author Contributions

Conceptualization, J.P. and G.A.; Methodology, J.P., R.A., B.J.H., B.A.K., P.J., and G.A.;Investigation, J.P., R.A., and B.J.H.; Resources, B.A.K., P.J., and G.A.; Writing – Original Draft, J.P.; Writing – Review & Editing, J.P., R.A., M.C.M., B.A.K., P.J., and G.A.; Visualization, J.P. and G.A.; Funding Acquisition, M.C.M., B.A.K., P.J., and G.A.; Supervision, B.A.K., P.J., and G.A.

## Acknowledgements

The authors thank Dr. Sukrit Singh for valuable discussion about the nature of mutual information as calculated by CARDS. This work was supported by National institutes of Health grants GM118817 (to G.A.), GM122979 (to P.J. and M.C.M.), and National Science Foundation grant MCB1817338 (to B.A.K.). Computational resources were provided by the NSF XSEDE Program ACI-1053575. The authors declare no conflict of interest.

