## Supplemental Figures for "Viral Packaging ATPases Utilize a Glutamate Switch to Couple ATPase Activity and DNA Translocation"

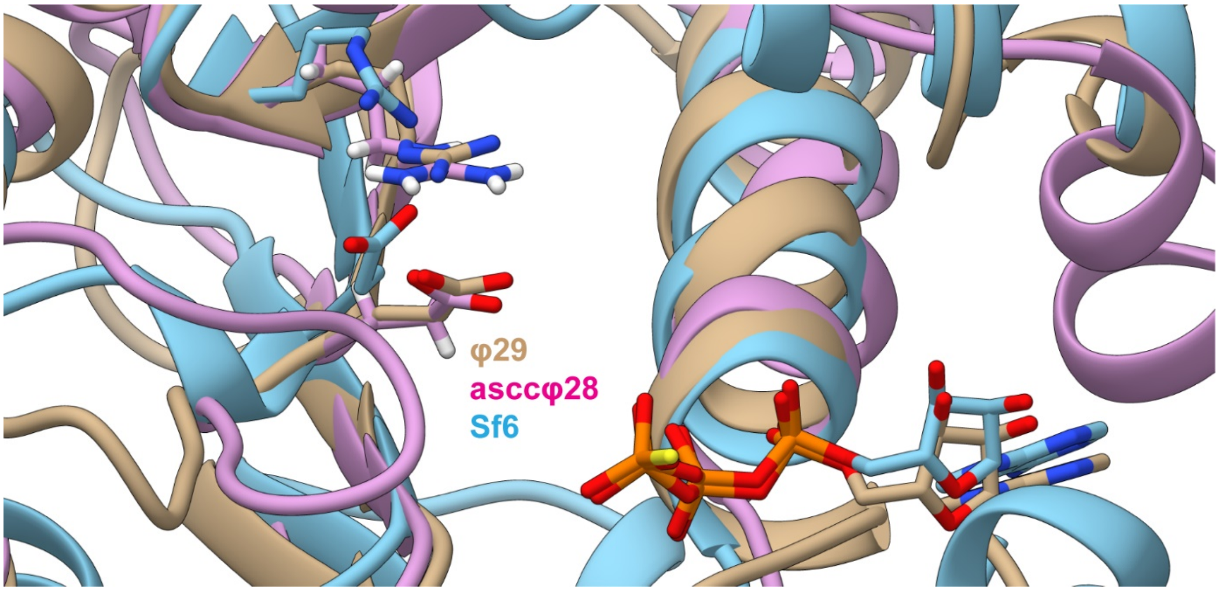


**Figure S1: Crystal structures of the φ29, asccφ28, and Sf6 packaging ATPases show that the catalytic glutamate residue points away from ATP (shown for clarity) and towards analogous arginines.**

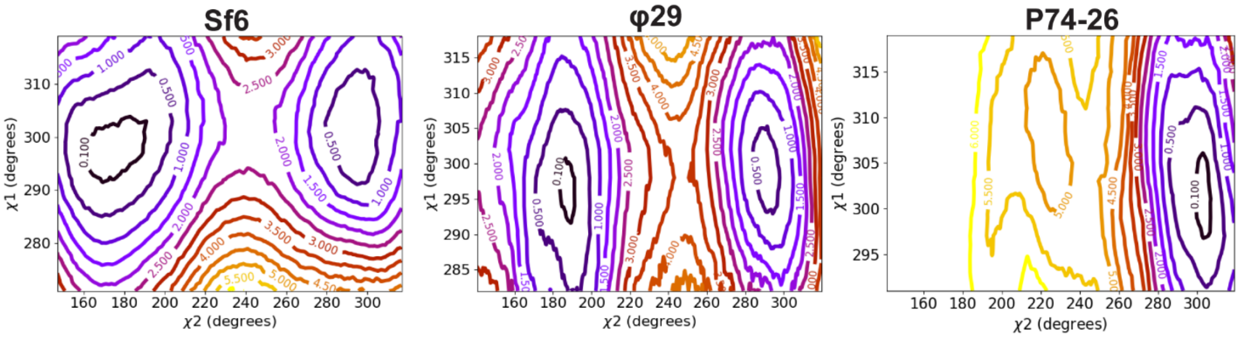


**Figure S2: Two-dimensional free-energy landscapes of apo conformations.** We find that the Sf6 and φ29 ATPases are stable in the inactive conformation, whereas the P74-26 ATPase has a metastability. This is attributed to differences in glutamate switch residues – Sf6 and φ29 use positively-charged arginine residues, whereas P74-26 uses a polar glutamine residue.

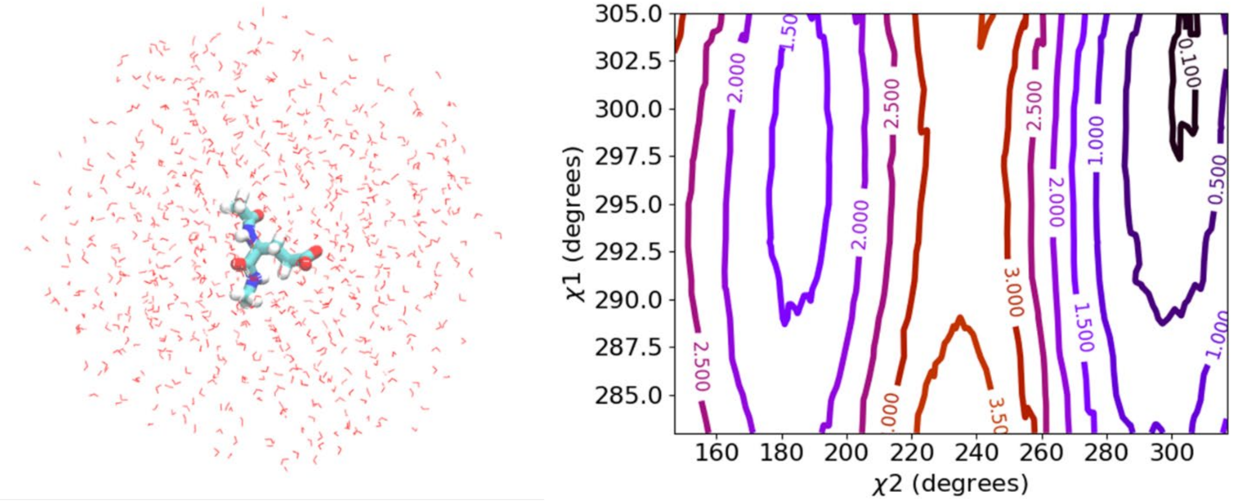


**Figure S3: Two-dimensional free-energy landscapes of a single glutamate residue in solution.** **(Left)** Schematic of simulation set up, with a single glutamate residue and We find that the energetic minimum for a single glutamate residue capped with an acetyl group on the N-terminus and n-methylamide on the C-terminus in a truncated octahedral periodic box. **(Right)** The two-dimensional free-energy landscape of the glutamate shows preference for the active pose. Thus, any preference for inactive pose found in enzymes is a result of interactions with other residues in the enzyme.

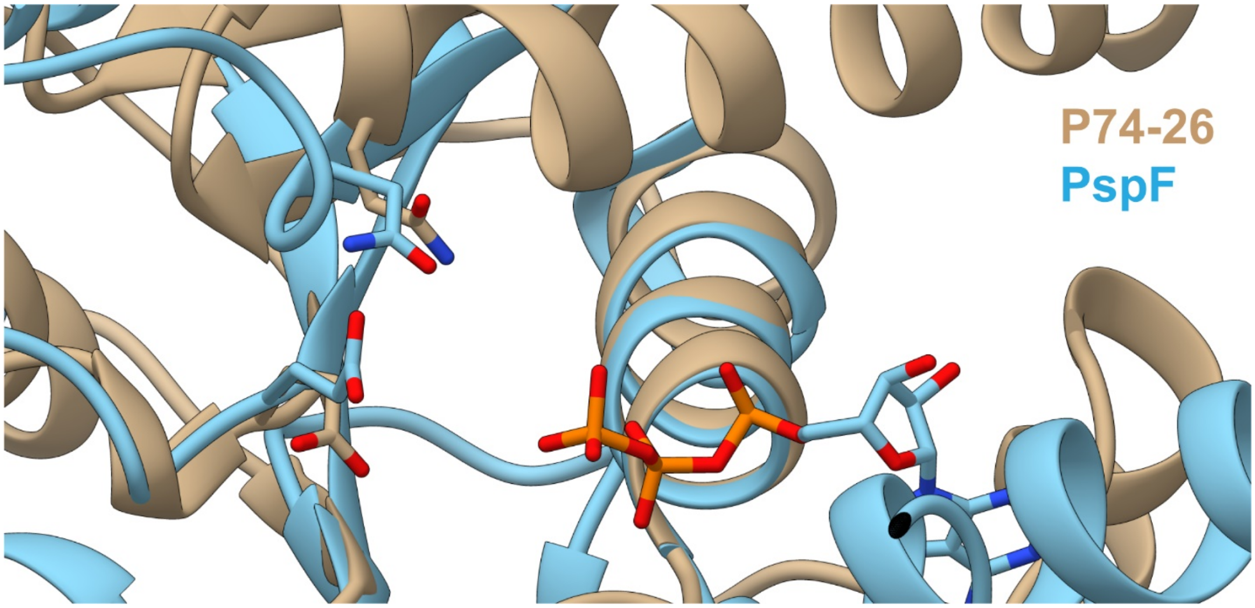


**Figure S4: Three-dimensional alignment suggests Gln72 is a glutamate switch in the P74-26 packaging ATPase.** The P74-26 packaging ATPase is aligned with the crystal structure of PspF bound with ATP (PDB: 2C96) based on their Walker A motifs. Upon alignment, P74-26 Gln72 is positioned similarly to PspF Asn64, which is one of the original glutamate switch residues identified. Thus, we considered that P74-26 Gln72 may also be a glutamate switch residue.

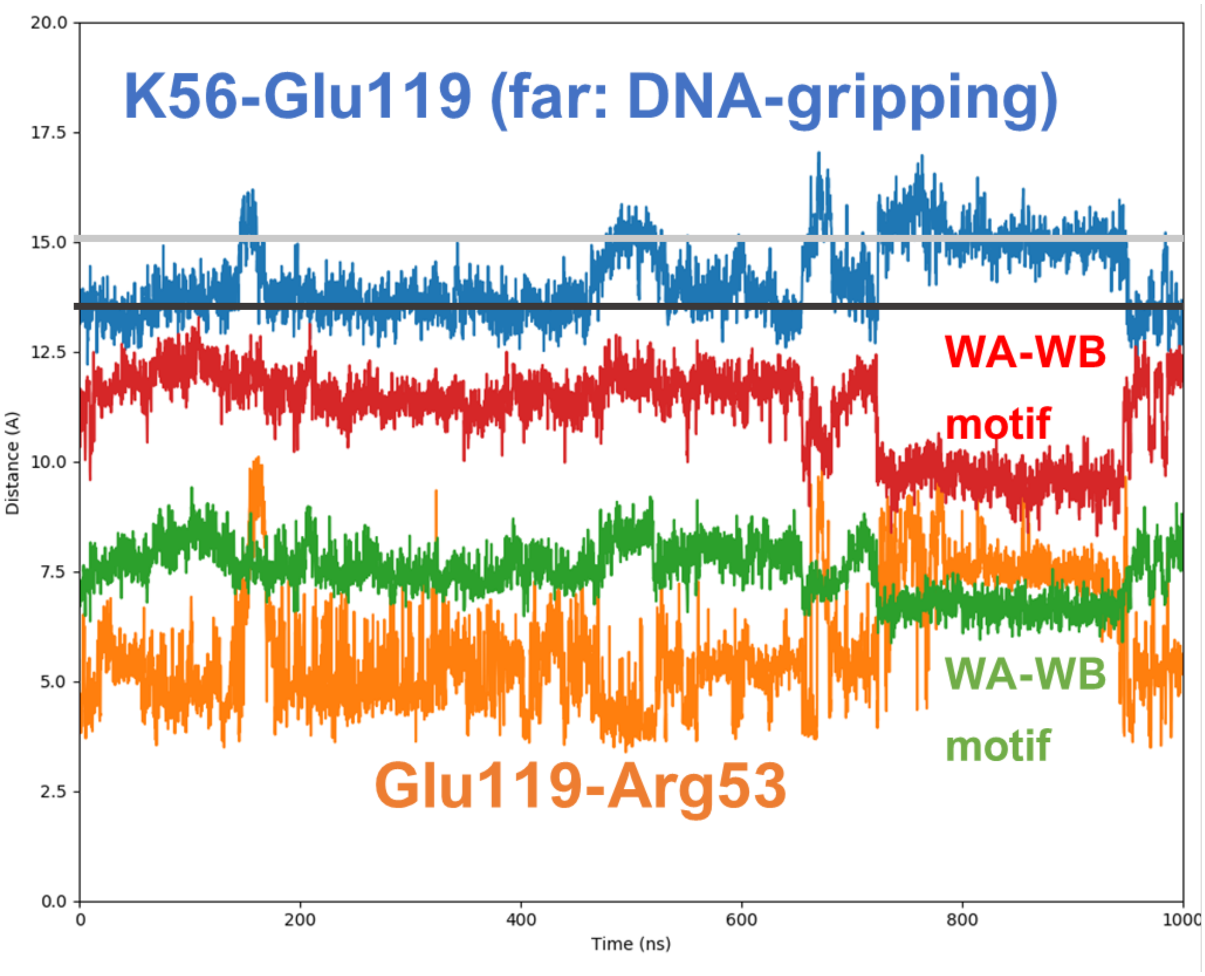


**Figure S5: Characterizing tight-binding in the φ29 packaging ATPase from equilibrium MD simulation.** Despite the φ29 crystal structure not containing the lid-subdomain, the tight-binding transition can still be characterized by the distance between Walker A (WA) and Walker B (WB) motif backbones. Upon tight-binding of the WA and WB motifs, the distance between the catalytic Glu119 and glutamate switch Arg53 increases, releasing the glutamate switch's hold of the catalytic glutamate. This release is accompanied by an increased distance between DNA-gripping Lys56 and Glu119 (from the black to the gray line), representing the donation of Kys56 farther into the pore to grip DNA. Umbrella sampling of the "tight-binding pose" starts from a frame taken between 800 and 950 ns, and umbrella sampling of the "pre-tight-binding" pose starts from a frame taken between 100 and 150 ns.

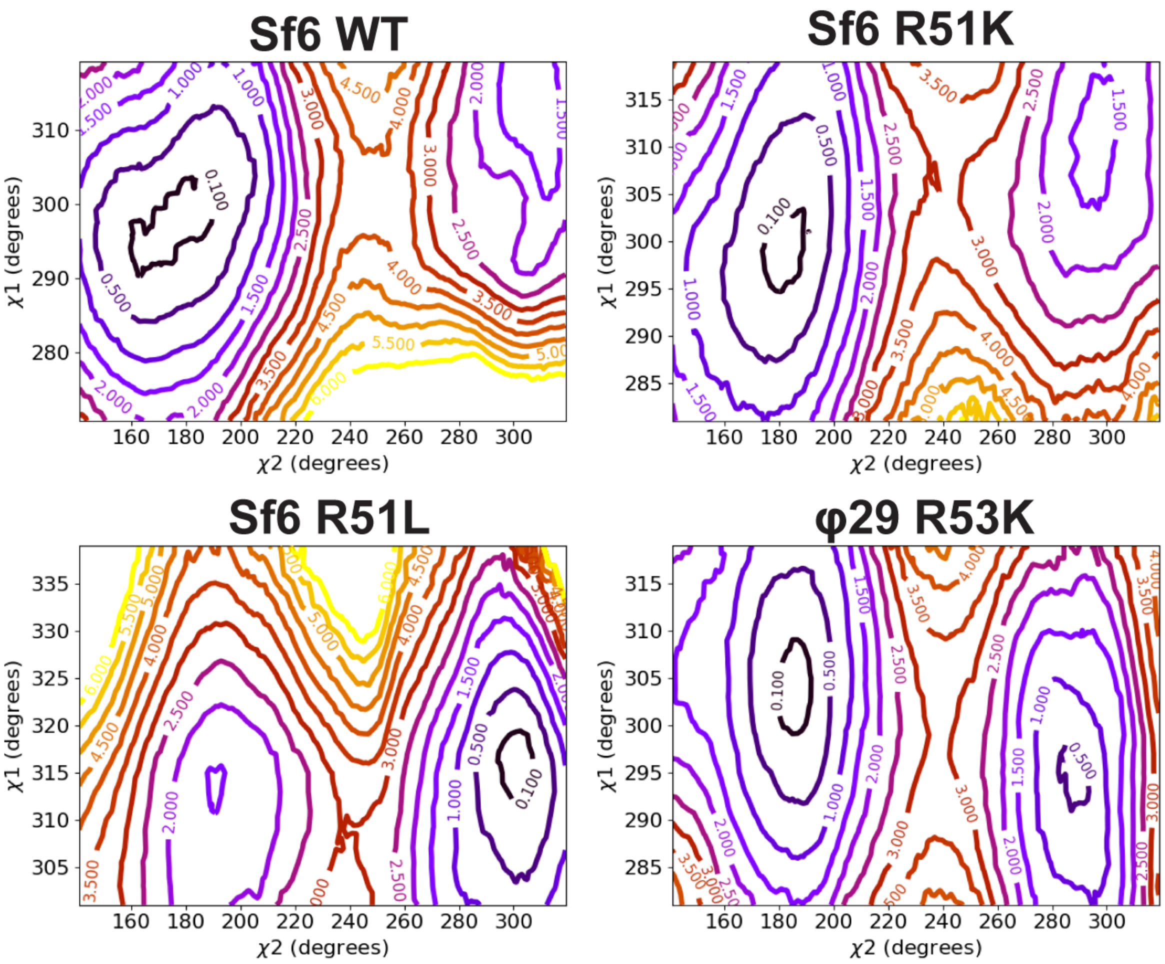


**Figure S6: Two-dimensional free-energy landscapes of mutants assign the role of the glutamate switch to a single residue.** We find that, in pre-tight-binding conformations, glutamate switch R→K does not affect preference for the inactive pose, whereas swapping R→L (see also main text **Fig. 3**) swaps preference from the inactive pose to the active pose. Thus, the positive charge on the conserved arginine is necessary to hold the catalytic glutamate residue inactive, and the arginine is a glutamate switch.

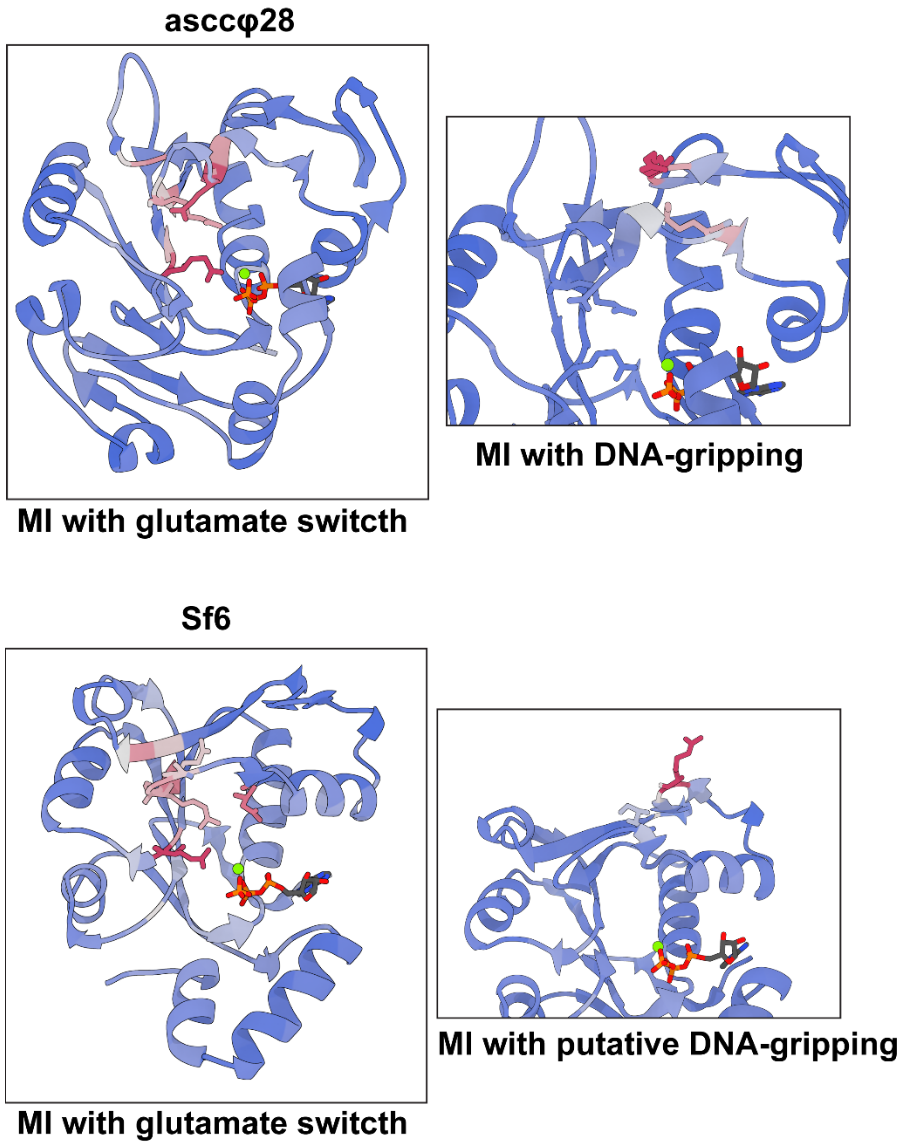


**Figure S7: Signaling pathways for asccφ28 and Sf6 predicted by CARDS mutual information.** For asccφ28 we find that the glutamate switch residue has high mutual information with both the catalytic glutamate and a second glutamate residue in the partitioning helical motif **(upper left panel)**, much like what we observe for φ29 (see main text **Fig. 5**). Like P74-26, (main text **Fig. 4** and **Fig. 5**), the partitioning helical motif can then push on a DNA-gripping arginine **(upper right panel)**, allowing the enzyme to position the DNA-gripping arginine in response to ATP-binding. For Sf6, we find that the glutamate switch has high mutual information with both the catalytic glutamate and an aspartate in the partitioning helical motif, allowing the enzyme to position this helix based upon ATP-binding. Though the DNA-gripping residues of the Sf6 ATPase are not experimentally verified, we note that there is a glutamine residue with high mutual information to the glutamate switch in the same position as φ29 Lys56, which is known to grip DNA **(lower left panel)**. An arginine analogous to the DNA-gripping residue in asccφ28 has relatively low mutual information with residues outside of its own motif, but does share information with a bulky hydrophobic residue in the partitioning helix situated between the glutamate switch and accompanying aspartate; this could indicate moving the helix acts sterically to position the DNA-gripping arginine **(lower right panel)**.
